# Antiviral *Wolbachia* strains associate with *Aedes aegypti* endoplasmic reticulum membranes and disturb host cell lipid distribution to restrict dengue virus replication

**DOI:** 10.1101/2023.09.14.557745

**Authors:** Robson K. Loterio, Ebony A. Monson, Rachel Templin, Jyotika T. Bruyne, Heather A. Flores, Jason M. Mackenzie, Georg Ramm, Karla J. Helbig, Cameron P. Simmons, Johanna E. Fraser

## Abstract

The insect endosymbiotic bacterium *Wolbachia pipientis* is being utilised as a biocontrol tool to reduce the incidence of *Aedes aegypti*-transmitted viral diseases like dengue. However, the precise mechanisms underpinning *Wolbachia*’s antiviral activity are not well defined. Here we generated a panel of *Ae. aegypti*-derived cell lines infected with antiviral strains *w*Mel and *w*AlbB or the non-antiviral strain *w*Pip to understand host cell morphological changes specifically induced by antiviral strains. Antiviral strains were frequently found to be entirely wrapped by the host endoplasmic reticulum (ER) membrane, while *w*Pip bacteria clustered separately in the host cell cytoplasm. ER-derived lipid droplets (LDs) increased in volume in *w*Mel-and *w*AlbB-infected cell lines and mosquito tissues compared to cells infected with *w*Pip or *Wolbachia*-free controls. Inhibition of fatty acid synthase (required for triacylglycerol biosynthesis) reduced LD formation and significantly restored ER-associated dengue virus replication in cells occupied by *w*Mel. Together, this suggests that antiviral *Wolbachia* strains may specifically alter the lipid composition of the ER to preclude the establishment of DENV replication complexes. Defining *Wolbachia*’s antiviral mechanisms will support the application and longevity of this effective biocontrol tool that is already being used at scale.

**Importance:** *Aedes aegypti* transmits a range of important human pathogenic viruses like dengue. However, infection of *Ae. aegypti* with the insect endosymbiotic bacterium, *Wolbachia*, reduces the risk of mosquito to human viral transmission. *Wolbachia* is being utilized at field sites across more than 13 countries to reduce the incidence of viruses like dengue, but it is not well understood how *Wolbachia* induces its antiviral effects. To examine this at the subcellular level, we compared how different strains of *Wolbachia* with varying antiviral strengths, associate with and modify host cell structures. Strongly antiviral strains were found to specifically associate with the host endoplasmic reticulum and induce striking impacts on host cell lipid distribution. Inhibiting *Wolbachia*-induced lipid redistribution partially restored dengue virus replication demonstrating this is a contributing role for *Wolbachia*’s antiviral activity. These findings provide new insights into how antiviral *Wolbachia* strains associate with and modify *Ae. aegypti* host cells.

## 1. Introduction

Arthropod-borne viruses (arboviruses) including dengue (DENV), Zika (ZIKV), chikungunya (CHIKV), and yellow fever virus (YFV), are primarily transmitted by female *Aedes aegypti* and, to a lesser extent, by female *Aedes albopictus* ^1–3^. The global spread of these viruses has dramatically increased in recent decades. This is largely due to factors associated with the geographic distribution of *Aedes* spp. such as climate change ^4–7^, globalization ^8,9^, urbanization ^9,10^, resistance to insecticides ^11^, and the lack of effective vector control strategies ^12^.

A promising biological vector control strategy that has emerged in the last decade employs *Wolbachia pipientis* to restrict mosquito-to-human transmission of *Ae. aegypti*-borne viruses ^13–17^. *Wolbachia* is an intracellular gram-negative endosymbiotic bacterium that infects a wide range of invertebrates, including arthropods and nematodes, but it is not naturally found in *Ae. aegypti* mosquitoes _18,19_. *Wolbachia* strains can be isolated from native hosts such as *Drosophila melanogaster* (*w*Mel) ^14,20^ and *Ae. albopictus* (*w*AlbB) ^15,21,22^, and then introduced into heterologous arthropods including *Ae. aegypti*. Strains such as *w*Mel and *w*AlbB induce an antiviral state in *Ae. aegypti* reducing the rate of infection, dissemination, and transmission of +RNA viruses such as DENV ^14,20,23^. *Wolbachia*’s exceptional ability to infect and alter the host germ line to facilitate their vertical transmission through the maternal lineage underpins the application of this bacterium as a biocontrol tool^14^. Additionally, many *Wolbachia* strains are capable of inducing cytoplasmic incompatibility (CI), a condition that gives *Wolbachia*-carrying females a reproductive advantage ^24,25^. These strains can be introgressed into wild-type *Ae. aegypti* populations to reduce their transmission potential. This approach has been shown to dramatically and significantly reduce the incidence of dengue in communities ^26–31^.

DENV is a member of the *Flaviviridae* family with a single positive-strand RNA genome that is surrounded by the virally encoded capsid protein in a host-derived lipid bilayer ^3,32^. The viral genome is released into the host cell cytosol after entering susceptible cells via endocytosis and is translated by host machinery at the rough endoplasmic reticulum ^33,34^. DENV induces drastic rearrangements of endoplasmic reticulum (ER) membranes to form viral replication complexes that are critically required to coordinate the multiple steps of viral replication, genome translation, and virion assembly ^33–39^. *Wolbachia*’s antiviral impacts are believed to be systemic in mosquitoes^40^, but also cell autonomous whereby *Wolbachia*-infected cells do not protect surrounding *Wolbachia*-free cells from viral infection ^41^. At the subcellular level, viral restriction is believed to occur early, before viral replication is effectively initiated ^42^, and low levels of progeny virus produced in the presence of *Wolbachia* have reduced infectivity ^43^. How *Wolbachia* induces these antiviral effects is not well understood, but a variety of hypotheses have been examined. These include competition between *Wolbachia* and viruses for nutrients and physical space within host cells ^13,44–48^, and altered expression of host pro-or antiviral genes ^49–52^, including priming of immune pathways ^23,53,54^. Implicating the mechanisms that drive the *Wolbachia*-induced antiviral state has been difficult due to technical limitations such as failure to genetically modify *Wolbachia* and to grow it axenically. However, new experimental models may help to advance our understanding of *Wolbachia*-induced host phenotypes. Specifically, we recently found that *Wolbachia* strain *w*Pip (derived from *Culex quinquefasciatus*) does not inhibit flavivirus replication, dissemination or transmission in *Ae. aegypti* ^23,55^. We hypothesized that pairwise comparisons between *Ae. aegypti* infected with *Wolbachia* strains that do or do not induce an antiviral state may facilitate the dissociation of *Wolbachia*’s antiviral effects from the general symbiont effects. Here we generated a unique panel of *Ae. aegypti*-derived cell lines infected with *w*Mel, *w*AlbB (the antiviral stains being utilized in *Ae. aegypti* biocontrol programs) or *w*Pip to identify the subcellular changes specifically induced by antiviral *Wolbachia* strains. We identify intimate *Wolbachia*-ER interactions and triacylglyceride biosynthesis for LD formation as key host cell modifications that contribute to the *Wolbachia*-induced antiviral state.

## 2. Results

### 2.1 Generation of a panel of Wolbachia-infected Ae. aegypti-derived cell lines

To examine the subcellular changes specifically induced by antiviral *Wolbachia* strains, we generated a panel of *Wolbachia*-infected cell lines using the immunocompetent *Ae. aegypti*-derived cell line (*Aag2*). *Aag2* cells infected with antiviral strain *w*Mel and the *w*Mel-cured line (*w*Mel.Tet-*Aag2*) have been described previously ^56^. *w*AlbB-*Aag2* and *w*Pip-*Aag2* cell lines were produced by stable infection of *w*Mel.Tet-*Aag2* using *Wolbachia* isolated from previously generated cell lines (RML12-*w*AlbB) or *Ae. aegypti* eggs (Rockefeller-*w*Pip). This was done to account for insect-specific flaviviruses known to infect *Aag2* cells but not *w*Mel-*Aag2* or *w*Mel.Tet-*Aag2* (herein referred to as *Wolbachia*-free-*Aag2*) ^57^.

To confirm that *w*AlbB and *w*Pip had antiviral and non-antiviral impacts in these cell lines, respectively, we evaluated DENV-2 replication in *Wolbachia*-free, *w*Mel-*Aag2*, *w*AlbB-*Aag2* and *w*Pip-*Aag2* cells. Consistent with our previous findings in *Ae. aegypti* ^23,55^, *w*Pip did not restrict DENV-2 replication or production of infectious virus compared to the *Wolbachia*-free line (Fig. 1A & B). Notably, *w*Pip was found to grow to a higher density than *w*Mel and *w*AlbB (average of 96 *Wolbachia* per cell, compared to 30 and 69 for *w*Mel and *w*AlbB, respectively) (Fig. 1A, in parentheses). By contrast, DENV-2 RNA copies and infectious virus were significantly reduced by *w*Mel’s potent antiviral activity (approximately 3 log_10_ reduction compared to *Wolbachia*-free). *w*AlbB also significantly inhibited DENV-2 replication and infectious virus in *Aag2* cell lines but the antiviral activity was weaker than *w*Mel (approximately 1 log_10_ reduction compared to *Wolbachia*-free). This confirms that the diverse antiviral phenotypes of *Wolbachia* strains previously demonstrated in *Ae. aegypti* mosquitoes ^20,22,23^, can be recapitulated in *Aag2* cell lines.

**Figure 1:**
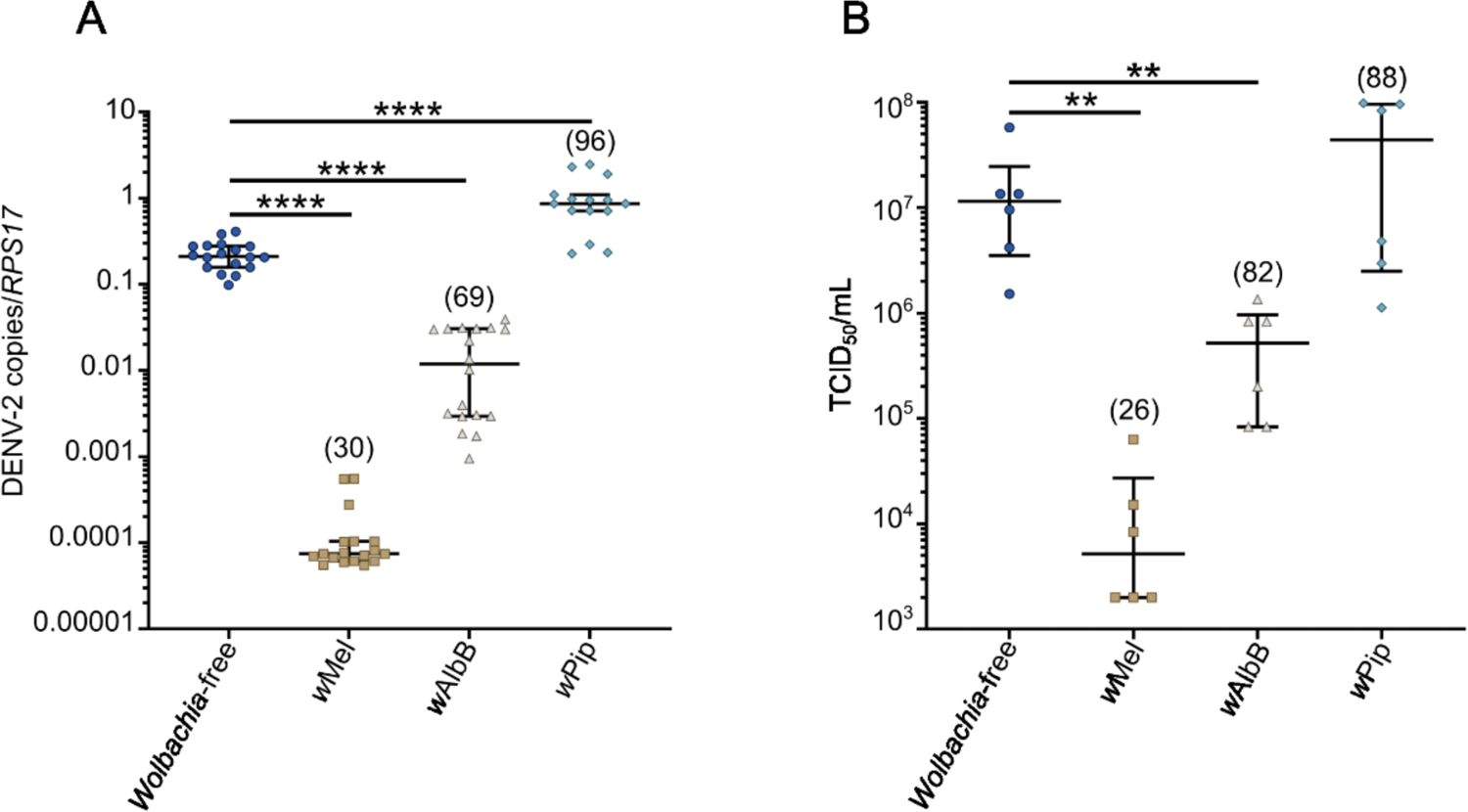
*w*Pip does not restrict DENV-2 replication in *Aag2* cells. *Ae. aegypti*-derived cell lines (*Aag2*) stably infected with *w*Mel, *w*AlbB, and *w*Pip strains were infected with DENV-2 at MOI 1 and compared to their matched *Wolbachia*-free line. Infectious virus and cellular RNA were collected from each cell line 7 days post-infection for analysis by TCID_50_ and qRT-PCR. The numbers in parentheses represent the average number of *Wolbachia* per cell (*Wolbachia 16S rRNA/RPS17*), determined in parallel wells for each independent experiment prior to DENV-2 inoculation. (A) Data are the median number of virus genome copies relative to the *RPS17* mosquito housekeeping gene ± IQR. (B) Data are the median infectious viral titres ± IQR determined by TCID_50_. Data are derived from at least 3 (A) or 2 (B) independent experiments performed with triplicate biological replicates each. Statistical analyses were performed using Mann-Whitney test, where ** p < 0.005, **** p < 0.0001.

### 2.2 Antiviral Wolbachia strains associate with host ER membranes

*Wolbachia* and viruses are both obligate intracellular residents of eukaryotic cells that rely on a variety of host structures and processes to complete their life cycles. Interestingly, *Wolbachia* titers and intracellular distribution have been shown to be influenced by the host’s genetic background ^46,58,59^. To gain insights into how *Wolbachia* strains may associate with *Ae. aegypti* cells to induce an antiviral phenotype, we assessed our panel of *Aag2* cell lines using transmission electron microscopy (TEM). Micrographs of the three *Wolbachia*-infected mosquito cell lines show each cell line is heavily infected with their respective *Wolbachia* strain (top row, Fig. 2A and Supplementary Fig. 1). *Wolbachia* appeared similar in size and shape to mitochondria but could be differentiated by a lower electron density and an absence of cristae. We observed that antiviral strains, *w*Mel and *w*AlbB, were frequently associated with host membranes that appeared to be the host endoplasmic reticulum (ER), while the non-antiviral strain *w*Pip was not (bottom row, Fig. 2A and Supplementary Fig. 1e-p). By scoring the closeness of interaction between *Wolbachia* and the ER a striking difference was evident between antiviral and non-antiviral strains: approximately 50% of *w*Mel and *w*AlbB bacteria were in close contact with ER membranes in *Aag2* cell lines. Of those, 25% were clearly surrounded by the membranes (Fig. 2B & C). This tight association was observed in our reconstructed tomograms of *w*Mel where the bacterium was found to be surrounded by the ER double membrane, and in some instances was wrapped multiple times (Fig. C – See Supplementary Movie 1 for the reconstructed tomogram). Meanwhile, over 75% of *w*Pip bacteria were not located near any intracellular membranes, and only 25% were situated adjacent to any part of the ER (Fig. 2B and Supplementary Fig. 1m-p). To further confirm that the membrane structures were indeed ER, we carried out live cell imaging using an ER tracker dye which binds to the sulfonylurea receptors of ATP-sensitive K^+^ channels present on ER membranes, and SYTO11 which preferentially stains *Wolbachia* DNA over eukaryote DNA ^58,60^. Confocal fluorescence imaging confirmed the antiviral strains *w*Mel and *w*AlbB exhibited substantial colocalization with ER membranes (white arrowheads, Fig. 2D), while *w*Pip did not, instead forming a massive cluster of cytosolic *Wolbachia* (Fig. 2D).

**Figure 2:**
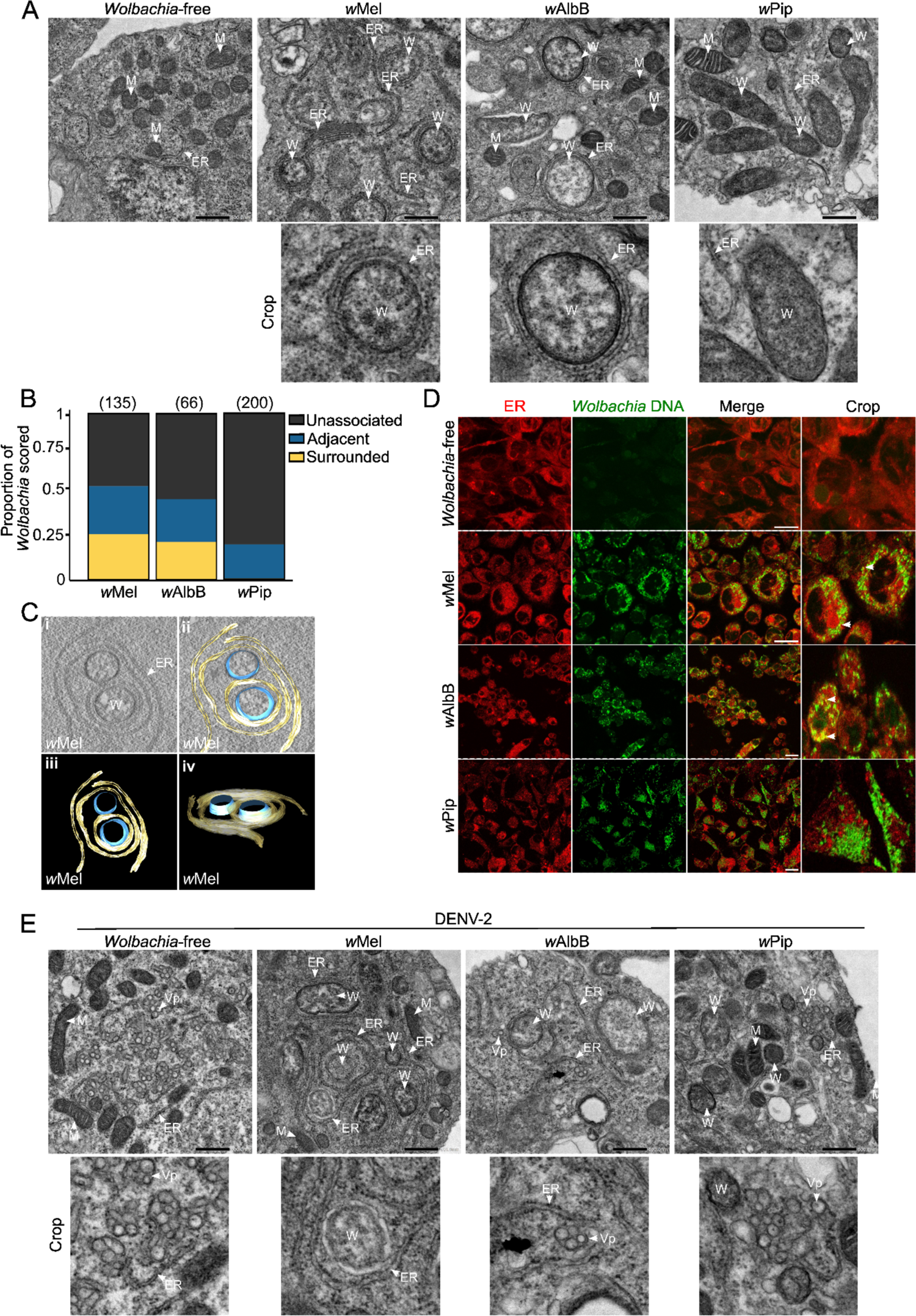
Antiviral *Wolbachia* strains associate with host endoplasmic reticulum membranes. (A) TEM micrographs of *Aag2* cell lines stably infected with *w*Mel, *w*AlbB, and *w*Pip show their intracellular distribution and association with ER membranes. Scale bar = 500nm. (B) *Wolbachia*-ER interactions were manually scored as unassociated, adjacent or surrounded from 2 independent experiments, where *Wolbachia* were counted from a minimum of 28 cells in total per line. The number of *Wolbachia* bacteria scored is in parentheses. (C) TEM tomography shows *w*Mel strain surrounded by ER membranes. See Supplementary Movie 1 for the reconstructed tomogram. (D) Live imaging of all *Aag2* cell lines stained with SYTO 11 (*Wolbachia* DNA - green) and ER tracker (red). Scale bar = 15μm. (E) TEM of all *Aag2* cell lines infected with DENV-2 at MOI 2 for 48 hours. Scale bar = 500nm. ER-Endoplasmic reticulum. W - *Wolbachia*. M-Mitochondria. Vp - Vesicle packets.

DENV and other flaviviruses significantly remodel ER membranes to form replication complexes ^32,33,35,36,61^. To determine whether the distinct ER interaction of antiviral *Wolbachia* strains precludes DENV-2 association with the host ER we infected each *Aag2* cell line with DENV-2 and performed TEM analyses. We observed numerous viral replication sites containing many viral vesicle packets (Vp) in the cytosol of the *Wolbachia*-free line (white arrowheads, Fig. 2E), consistent with the high viral titers determined in this cell line (Fig. 1). We found no evidence of virus replication in any of the *w*Mel-*Aag2* cell line micrographs examined, and few and smaller Vp were observed in the *w*AlbB-*Aag2* cell line (white arrowheads, Fig. 2E). Further consistent with our data in Fig. 1, *w*Pip showed numerous Vp in the cytosol, similar to the *Wolbachia*-free control (white arrowheads, Fig. 2E). Together this dataset provides the first insight into how antiviral *Wolbachia* strains differentially associate with *Ae. aegypti* ER membranes and prevent formation of typical DENV-2 replication complexes.

### 2.3 Antiviral Wolbachia strains increase lipid droplet formation in Aag2 **cells**

It has previously been reported that the ER-derived organelle, lipid droplets (LDs) are upregulated following DENV infection, contributing to the cellular antiviral response in both mammalian and *Ae. aegypti*-derived cell lines ^62,63^. Since the ER provides most of the constituent molecules for LDs ^64^, we next hypothesised that the association of antiviral *Wolbachia* strains with host ER membranes could further manipulate other intracellular lipid sources.

To examine intracellular LDs, we stained all *Aag2* cell lines with BODIPY 493/503 and analyzed the number and size of LDs per cell. Notably, compared to the *Wolbachia*-free line, all *Wolbachia*-infected lines induced LD accumulation (Fig. 3A & B). Interestingly, *w*Mel induced the accumulation of a similar number of LD’s per cell as *w*Pip (27 and 30.2 per cell, respectively), while *w*AlbB induced near twice the amount (50.5 per cell). However, *w*Mel-induced LDs were significantly larger than those induced by *w*AlbB and *w*Pip (791.5, 417.5, and 365.5 nm respectively) (Fig. 3B). This implies that antiviral *Wolbachia* strains might boost the total volume of LDs per cell.

**Figure 3:**
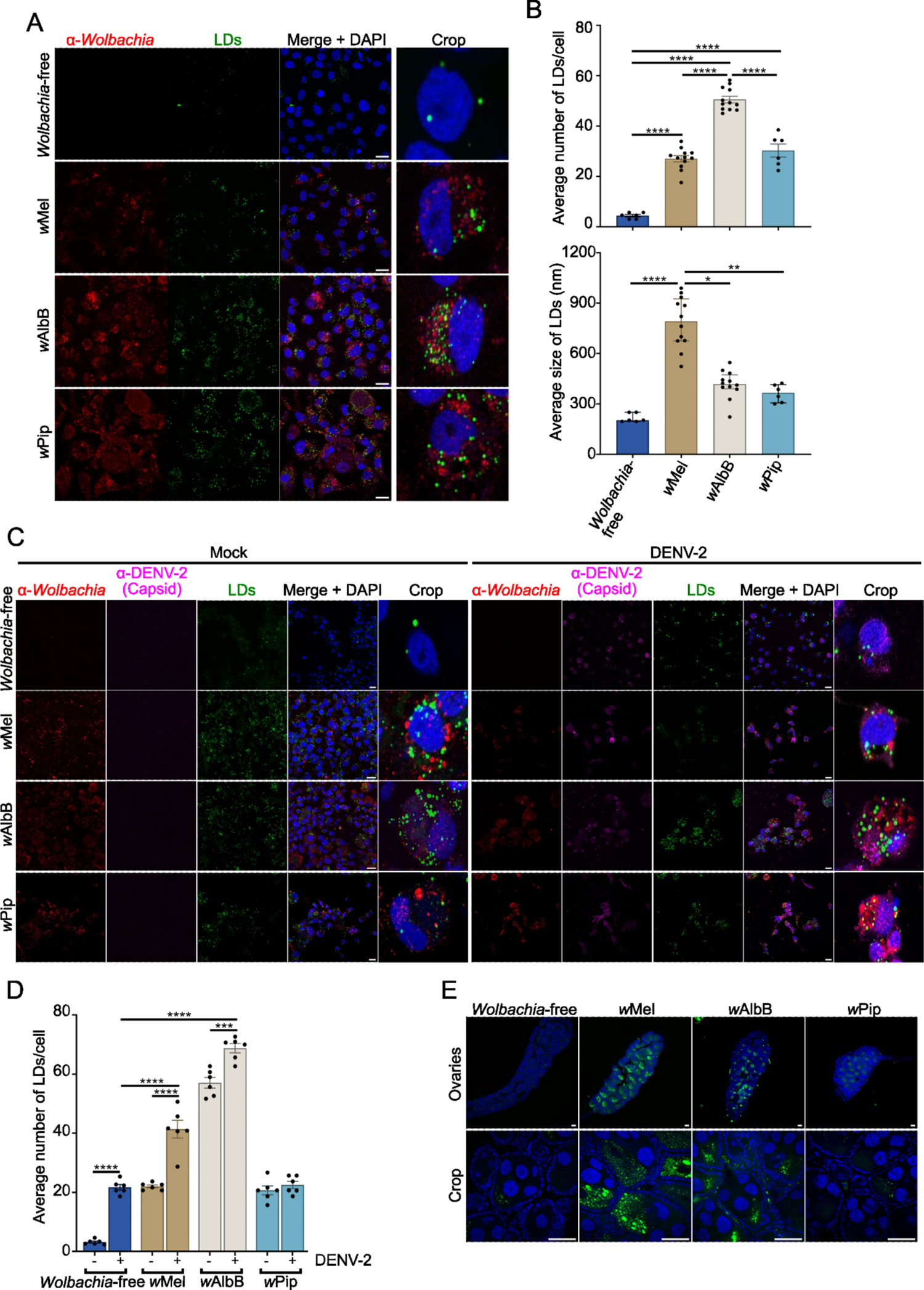
Antiviral *Wolbachia* strains increase lipid droplet formation in mosquito cells. (A-D) *Aag2* cells stably infected with *Wolbachia* strains *w*Mel, *w*AlbB, or *w*Pip, or *Wolbachia*-free cells were mock-infected or DENV-2 infected at MOI 1 for 24 hours. (A and C) All cells were stained with BODIPY (493/503) to visualize LDs (green) and DAPI to visualize the cell nuclei (blue). *Wolbachia* was detected with an α-WSP antibody (red) and 4G2 hybridoma fluid against flavivirus group E antigen for DENV-2 staining (magenta). Scale bars = 15μm. (B) LD numbers and sizes were analyzed using Fiji software. Data are the mean LD number ±SEM (top) or median LD size ±IQR (bottom) from at least 6 fields of view representing at least 80 cells in total where each data point represents the mean LD number/size from a single field of view. Statistical analyses were performed by One-way ANOVA with Tukey’s correction for multiple comparisons (top graph) or Kruskal-Wallis test with Dunn’s correction for multiple comparisons (bottom graph). (D) LD numbers were analyzed using Fiji software. Data are the mean LD number ±SEM from at least 6 fields of view representing at least 80 cells in total where each data point represents the mean LD number from a single field of view. Statistical analyses were performed by Two-way ANOVA with Tukey’s correction for multiple comparisons, * p < 0.05, ** p < 0.005, *** p < 0.0005, **** p < 0.0001. (E) Ovaries were dissected from female mosquitoes 5-7 days post-emergence and stained with BODIPY (green) and DAPI (blue). Scale bars = 50μm. All slides were imaged as 3-dimensional z-stacks and 2D images generated by Maximum Intensity Projection (MIP) using Fiji software.

Lipid biosynthesis and degradation play a role in several stages of DENV infection, including viral replication, assembly, and energy supply ^35,65–67^. Similarly, *Wolbachia* proliferation is tightly associated with changes in the host lipidome ^68–70^. Next, we investigated whether DENV-2 infection affected LD accumulation in the presence of each *Wolbachia* strain. In line with previous findings ^62^, DENV-2 induced the accumulation of LDs in *Wolbachia*-free cells (Fig. 3C & D). Interestingly, DENV-2 infection caused a further accumulation of LDs in *w*Mel-*Aag2* and *w*AlbB-*Aag2*, but not in *w*Pip-*Aag2* (Fig. 3D). Furthermore, the LD accumulation observed in both cell lines with antiviral *Wolbachia* strains after DENV-2 infection was significantly higher than the LD accumulation in *Wolbachia*-free cells infected with DENV-2 (Fig. 3D). By contrast, we measured similar levels of LDs in *w*Pip-and *Wolbachia*-free *Aag2* cells infected with DENV-2 (22.5 and 21.7 per cell, respectively) (Fig. 3D), indicating an association between high LD numbers and restriction of DENV-2 replication.

To investigate LD accumulation induced by different *Wolbachia* strains *in vivo*, we dissected female *Ae. aegypti* infected with *w*Mel, *w*AlbB or *w*Pip. We selected ovarian tissue since this tissue is rich in *Wolbachia* for all strains of interest (Supplementary Fig. 2) ^23^. Ovaries were stained with BODIPY 493/503 for LDs and DAPI to demarcate nuclei. Supporting our *in vitro* findings, LDs were most prevalent in the presence of antiviral *Wolbachia* strains compared to the *Wolbachia-*free control (Fig. 3E). Taken together, our findings demonstrate that changes in intracellular lipid storage are key features of mosquito cells infected with antiviral *Wolbachia* strains.

### 2.4 Intracellular redistribution of lipids contributes to DENV-2 restriction in ***w*Mel-*Aag2* cells**

We next hypothesized that *Wolbachia*-induced LD formation may be required to induce an antiviral state. A variety of small molecule inhibitors targeting mammalian enzymes involved in LD formation were applied to *w*Mel-and *Wolbachia*-free-*Aag2* cells to identify inhibitors that were effective against *Ae. aegypt*i orthologs. The DGAT 1 inhibitor (T863) and DGAT 2 inhibitor (PF-06424439) did not reduce LD formation in these cells, but the fatty acid synthase (FAS) inhibitor C75 did (Supplementary Fig. 3A & B). FAS is an enzyme that catalyzes the second step of the *de novo* triacylglycerol synthesis pathway contributing to lipid storage (LD formation) and membrane synthesis ^71,72^.

We therefore compared DENV-2 replication and infectious virus production in *Wolbachia*-free and *w*Mel-*Aag2* cell lines treated with C75. While C75 treatment reduced LD numbers in both cell lines DENV-2 replication was significantly reduced only in *Wolbachia*-free cells (by ∼1 log_10_), consistent with previous reports of LDs playing a key role in DENV replication ^67^ (Fig. 4A, B & C). By contrast, C75 treatment of the *w*Mel-*Aag2* cell line partially restored DENV-2 replication, with intracellular DENV-2 and infectious virus ∼1 log_10_ higher than in untreated *w*Mel-*Aag2* cells. Notably, C75 treatment of *w*Mel-*Aag2* cell lines did not reduce *Wolbachia* density (Fig. 4A & B, in parentheses), indicating that FAS and *w*Mel-induced LDs are not directly required for active *Wolbachia* replication and maintenance. However, it is possible that prolonged periods of LD depletion could affect *Wolbachia* survival.

**Figure 4.**
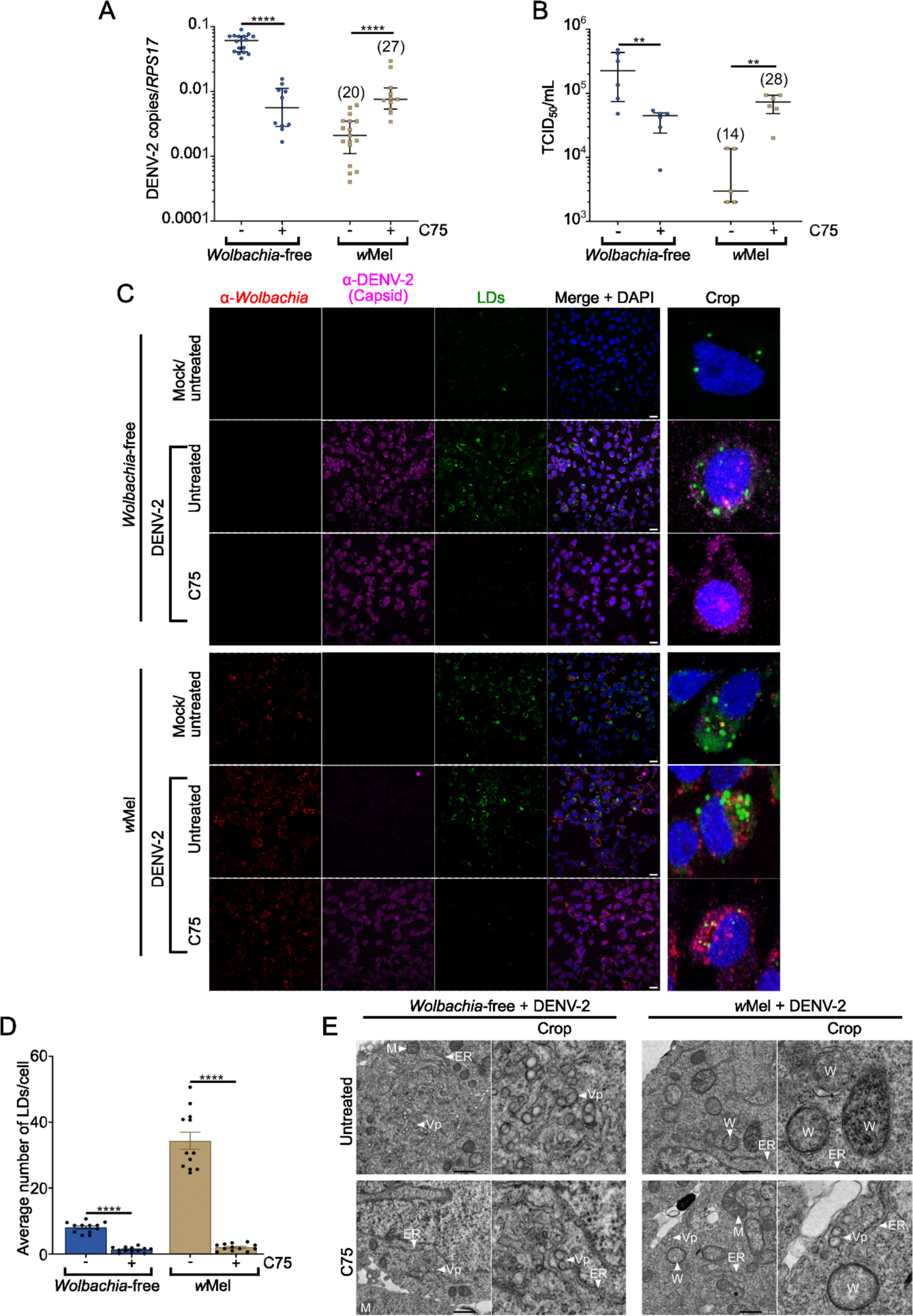
Disrupting *Wolbachia*-induced lipid droplets facilitates viral replication in *w*Mel-*Aag2* cell line. *Wolbachia*-free and *w*Mel-*Aag2* cell lines were mock-infected or DENV-2 infected at MOI 1 for 24 hours. Prior to DENV-2 infection cells were pre-treated with C75 for 24 hours and C75 was maintained during the time of infection. (A-B) Cell-associated RNA and virus supernatant were collected from each cell line 24 hours post-DENV-2-infection for analysis by qRT-PCR (A) and TCID_50_ (B). Data are the median number of viral genome copies ±IQR relative to the *RPS17* mosquito housekeeping gene (A) or the median TCID_50_/ml ±IQR (B). The numbers in parentheses represent the average number of *Wolbachia* per cell (*Wolbachia 16S rRNA/RPS17*), determined in parallel wells for each independent experiment prior to DENV-2 inoculation. Data are representative of at least 3 (A) or 2 independent experiments (B) with triplicate biological replicates each. Statistical analyses were performed using Mann-Whitney test, where ** p < 0.001 and **** p < 0.0001. (C) *Wolbachia*-free (top panel) and *w*Mel-*Aag2* cell lines (bottom panel) were stained with BODIPY (green) and DAPI (blue). *Wolbachia* was detected with an α-WSP antibody (red) and an 4G2 hybridoma fluid against flavivirus group E antigen for DENV-2 staining (magenta). Slides were imaged as 3-dimensional z-stacks and 2D images generated by MIP using Fiji software. Scale bars = 15μm. (D) LD numbers were quantified after 24 hours post-DENV-2 infection at MOI 1. LD numbers were analyzed using Fiji software. Data are the mean LD number ±SEM from at least 6 fields of view representing at least 80 cells in total, where each data point represents the mean LD number from a single field of view. Statistical analyses were performed by an unpaired two-tailed Student’s *t*-test, where **** p < 0.0001. (E) TEM of *Wolbachia*-free and *w*Mel-*Aag2* cell lines mock-infected or DENV-2 infected at MOI 1 for 24 hours and treated with C75 as previously described. Scale bar = 500nm. ER-Endoplasmic reticulum. W-*Wolbachia*. M-Mitochondria. Vp-Vesicle packets.

Finally, TEM micrographs of both *Wolbachia*-free and *w*Mel-*Aag2* cell lines treated with C75 were examined to verify whether DENV-2 replication could now be supported in cells near *w*Mel bacterium. We observed a reduction in visible cytosolic Vps in the *Wolbachia*-free cell line, while, for the first time, we were able to find evidence of virus replication in cells already occupied by *w*Mel (Fig. 4E). Together, these data show that blocking the *de novo* lipogenesis of triacylglycerol significantly compromises *w*Mel-induced LD accumulation, and consequently, its antiviral activity in *Ae. aegypti* cells.

## Discussion

*Wolbachia* has been implemented as a biocontrol tool in cities in Oceania, Asia, and Latin America. Several epidemiological studies have now demonstrated the efficacy of *Wolbachia*-introgression in reducing the burden of mosquito-borne diseases in communities ^26–31^ and the Vector Control Advisory Group to the World Health Organization has endorsed its public health value ^73^. To best support the longevity of this method, and to predict the emergence of *Wolbachia*-resistant viruses, it is critical that we understand the mechanisms that underpin the antiviral activity of *Wolbachia*.

Given that *Wolbachia* strains are classified into major phylogenetic lineages known as supergroups ^74^, we can strategically compare host modifications induced by the antiviral strain *w*AlbB and the non-antiviral strain *w*Pip because they both belong to supergroup B. Additionally, both strains are adapted to mosquito host species ^74,75^. *w*Mel, which belongs to supergroup A ^74^, has a well-characterized antiviral activity and comparisons with *w*AlbB enable us to determine whether antiviral strains from different supergroups employ similar strategies to induce the antiviral state in *Ae. aegypti*. In our cell culture models, we observed that supergroup B strains, *w*AlbB and *w*Pip, grew to substantially higher densities (>80 *Wolbachia*/cell) than the distantly related *w*Mel (<30 *Wolbachia*/cell). This is consistent with somatic tissues in mosquitoes where it has been reported that *w*AlbB and *w*Pip generally reside at higher levels than *w*Mel ^20,23,55^. Thus, our *in vitro* findings support previous research and show that *Wolbachia* density does not necessarily determine a strain’s antiviral activity.

Recent investigations have demonstrated that *w*Mel resides nearby ER and Golgi organelles in its native host, *D. melanogaster* ^46,47,58^. Here, we remarkably observed that only antiviral *Wolbachia* strains associate closely with the ER network in *Aag2* cell lines. Non-antiviral strain *w*Pip instead formed a massive cytoplasmatic cluster with no apparent interaction with other organelles. TEM analyses revealed *w*Mel and *w*AlbB were individually surrounded by ER membranes at a high frequency. This may be strategic by these strains; for example, the *w*Mel genome lacks pathways for metabolizing some membrane components, relying on its host for many of the materials required for their membrane formation ^76–78^. Since *w*Pip does not demand this close association for its active replication, perhaps it has alternate ways of acquiring nutrients or has a more complete intrinsic metabolic capacity. Comparative genomic studies between related strains *w*AlbB and wPip may provide insight into this. Interestingly, based on our TEM micrographs, even in *Aag2* cell lines highly infected with *w*AlbB or *w*Pip strains (>80 *Wolbachia*/cell), we did not see any morphological evidence of increased ER activity linked to ER stress, such as enlarged tubules and cisternae, nor subcellular redistribution of this organelle as previously described for *w*Mel in *D. melanogaster* ^46,47^.

DENV and other flaviviruses considerably remodel ER membranes for their own replication ^32,33,35,36,61^. While it is interesting that only the antiviral *Wolbachia* strains were found to be substantially associated with this organelle, our TEM analyses did not indicate any spatial competition between antiviral *Wolbachia* strains and DENV-2 for ER membranes. Indeed, ER membranes are prolific in eukaryotic cells and we observed large regions of *Aag2* cells that had ER membranes free from *Wolbachia*, with space to hypothetically support formation of viral replication complexes. Instead, we hypothesize that antiviral *Wolbachia* strains may biochemically alter the composition of the ER membranes in *Aag2* cells by eliciting the formation of LDs. LDs may then have further direct or indirect roles in mediating viral restriction.

LDs are thought to have conflicting roles in viral infection: many reports have demonstrated that enveloped viruses usurp LDs and alter lipidomic profiles of host cells to enhance their viral life cycles, with lipids accounting for 20-30% of the weight of the virion ^62,67,79–83^. DENV infection, for example, increases LD formation ^62^, recruiting FAS to the virus replication site ^65^, and its capsid protein accumulates on the surface of LDs, facilitating viral replication by providing a platform for nucleocapsid formation during encapsidation ^67^. By contrast, LDs have been shown to have a critical role in supporting innate immune signaling pathways, potentially acting as a platform to augment and coordinate the cell’s antiviral response ^62,63,84,85^. Since recent studies have suggested that *Wolbachia*’s antiviral activity towards DENV is not driven by innate immune priming ^23,53^ it remains unclear what role *Wolbachia*-induced LDs may have in viral restriction.

In this study we noticed distinct LD profiles in *Aag2* single-infected with *Wolbachia* strains. Antiviral *Wolbachia* strains induced a higher volume of LDs per cell than *w*Pip. Additionally, we found that *w*Mel-and *w*AlbB-*Aag2* cell lines responded to DENV-2 infection by further accumulating LDs, which did not occur in *w*Pip-*Aag2* cell line. Based on these findings, it appears that DENV-2 replication in *Aag2* cell lines requires a preferred LD accumulation threshold that, if crossed, e.g. in the presence of antiviral *Wolbachia* strains, no longer supports DENV-2 replication. Previously, *Aag2* cells infected with *w*MelPop (a supergroup A pathogenic antiviral *Wolbachia* strain from *D. melanogaster*) were shown to have LDs enriched in esterified cholesterol. DENV-2 replication was partially rescued in these cells by dispersing localized cholesterol using the drug 2-hydroxypropyl-β-cyclodextrin. However, this phenotype was not reproducible in *Ae. albopictus*-derived cells, *Aa23*, infected with another supergroup A antiviral *Wolbachia* strain, *w*Au ^86,87^. Therefore, assessment of all antiviral strains using a common approach will be important in the future to determine whether some antiviral strains utilize different antiviral mechanisms in *Ae. aegypti*.

Meanwhile, Manokaran and colleagues discovered that acyl-carnitines (intermediate molecules that transport activated fatty acids, FA-CoA, from the cytoplasm to the mitochondria) are downregulated in the presence of *w*Mel, re-directing lipid sources from β-oxidation, and negatively affecting ATP production, which is required for efficient DENV-1 and Zika virus replication ^48^. All these findings support our hypothesis that antiviral *Wolbachia* strains relocalize essential lipid classes and limit their availability to sustain DENV replication.

C75 inhibition of LD formation has previously been shown to impact DENV-2 RNA amplification in the absence of *Wolbachia*. This is thought to be because C75 inhibits association of the capsid protein with host LDs ^67,88^. Here, C75 treatment of *w*Mel-*Aag2* cells abrogated LD formation and instead supported an increase in intracellular and infectious titers of DENV-2, to levels comparable with C75-treated *Wolbachia*-free-*Aag2* cells. This partial restoration of DENV-2 replication in *w*Mel-*Aag2*s is consistent with excess LD’s induced by *w*Mel having an antiviral role, while also supporting a pro-viral role for moderate levels of LDs ^67,88^. That is, complete abrogation of LD formation as we observed in both *w*Mel-and *Wolbachia*-free *Aag2* cells treated with C75, is partially restrictive to DENV.

It is important to acknowledge that the C75 target, FAS, does not directly drive formation of LDs, and instead catalysis synthesis of triacylglycerols. Excess free fatty acids are converted into neutral lipids and stored in cytosolic LDs. Primarily, FAS synthesizes palmitate from acetyl-CoA and malonyl-CoA in the presence of NADPH_72_. Thus, although FAS inhibition was associated with reduced LD formation, we cannot exclude the possibility that other FAS-mediated changes in fatty acid homeostasis are responsible for the partial restoration of DENV-2 replication in *w*Mel-*Aag2* cells.

One impact of reducing LD formation is the accumulation of lipids at ER membranes, especially precursors of triacylglycerols and sterol esters (such as cholesterol), which constitute most of the structural core of LDs ^89,90^. Within the several classes of lipids, sphingolipids and sterols are essential in determining membrane flexibility and stability ^91,92^. Therefore, an adequate membrane-lipid composition in the ER is critical for membrane rearrangement and assembly and function of +RNA virus replication complexes ^32,33,35,36,61^. Thus, reduction of LD formation following C75 treatment might facilitate DENV-2 acquisition of lipid classes in the ER that may not normally be available in the presence of antiviral *Wolbachia* strains. Moreover, dysregulation of LDs could modify not only the composition of the ER membranes but also the composition of the LDs, both of which are required for DENV replication. In reality, *Wolbachia*’s redistribution of lipids within cells may mask intracellular competition for lipids. That is, differential localization of specific lipid classes that prevent effective viral replication on ER membranes may not have been detected in past lipidomic studies if the overall prevalence of each lipid class remains the same ^48,70^. The mechanism by which antiviral *Wolbachia* strains remove lipids from ER membranes and store them as LDs must therefore be further examined.

These results provide the first detailed analyses of how antiviral *Wolbachia* strains associate with and modify *Ae. aegypti* cell organelles to restrict viral replication. We identify two striking phenotypes induced only by antiviral strains, including intimate association with host ER membranes, and induction of bigger and/or more numerous LDs. Our data support a role for *Wolbachia*-induced lipid redistribution in restriction of DENV-2 replication.

## MATERIAL & METHODS

### Cell line generation and maintenance

*w*Mel-*Aedes aegypti-*derived *(Aag2*) cell line and the *Wolbachia*-free *w*Mel.Tet*-Aag2* cell lines have been described previously ^56^. Briefly, *w*Mel.Tet*-Aag2* was generated by treating *w*Mel-*Aag2* cell line with 10 µg/mL tetracycline for 3 successive passages, to cure the line of *w*Mel infection. This *Wolbachia*-free-*Aag2* cell line was then further used to generate *w*AlbB-*Aag2* and *w*Pip-*Aag2* cell lines. *w*AlbB strain was purified from the *Ae. albopictus*-derived cell line, RML-12, stably infected with *w*AlbB and infection of *Wolbachia*-free-*Aag2* cells was done using the shell vial technique as described previously ^59^.

To generate *w*Pip-*Aag2*, *w*Pip was extracted from infected *Ae. aegypti* eggs_93_. In an Eppendorf tube, 1000-2000 2-5-day old eggs were rinsed 3-5 times in 1mL distilled water. The eggs were passed through a 100 µm mesh sieve and placed in a new Eppendorf tube containing 1 mL 80% v/v ethanol. Eggs were sterilized by 3-5 washes in 80% v/v ethanol then rinsed three times in distilled water, sieved, and transferred to a new Eppendorf tube containing 500 µL SPG buffer (218 mM sucrose, 3.8 mM KH_2_PO_4_, 7.2 mM K_2_HPO_4_, 4.9 mM L-glutamate at pH 7.4). Eggs were rinsed three times in SPG buffer then homogenized in 300-500 µL SPG buffer using a plastic pestle. The homogenate was spun to remove debris, and the supernatant was collected in a fresh tube. This was repeated three to five times until the supernatant became clear. A pool of supernatant samples was created. Under sterile conditions, the *w*Pip-containing supernatant was then passed through a 5 µm syringe filter followed by a 2.7 µm syringe filter. The filtrate was then centrifuged at 12,000 g to pellet *Wolbachia*. Extracted *w*Pip pellet was finally resuspended in ∼300 µL of sterile SPG buffer. *Wolbachia*-free-*Aag2* cell line was then infected with the purified *w*Pip using the shell vial technique ^59^.

All *Aag2* cell lines (hereafter referred as *Wolbachia*-free-*Aag2*, *w*Mel-*Aag2*, *w*AlbB-*Aag2*, and *w*Pip-*Aag2* cell lines) were routinely cultured at 26°C in maintenance media consisting in 1:1 Schneider’s Drosophila (Gibco)/Mitsuhashi and Maramorosch (CaCl_2_ 0.151 g/L, MgCl_2_ 0.047 g/L, KCl 0.2 g/L, NaCl 7 g/L, NaH_2_PO_4_ 0.174 g/L, D(+)-glucose 4g/L, Yeast extract 5 g/L, Lactalbumin hydrolysate 6.5 g/L, and NaHCO_3_ 0.12g/L at pH 6.9) medium supplemented with 10% fetal bovine serum (FBS – Gibco).

C6/36 cell lines of *Aedes albopictus* origin were supplied by the American Type Culture Collection (ATCC) were routinely cultured at 28°C with 5% CO_2_ in RPMI 1640 medium (Gibco) with GlutaMAX^TM^ Supplement containing 10% FBS.

### Wolbachia Density

*Wolbachia* density was routinely monitored in each cell line. To determine the average number of *Wolbachia* per cell we used qPCR to measure the relative abundance of the conserved *Wolbachia 16S rRNA* gene to that of the single-copy mosquito house-keeping gene *RPS17* gene. *Wolbachia*-free cells and *w*Mel-, *w*AlbB-and *w*Pip-*Aag2* cell lines were lysed with squash extraction buffer (10 mM Tris Buffer, 1 mM ethylenediamine tetraacetic acid [EDTA], 50 mM NaCl in ultrapure water, 1:50 Proteinase K). Cell lysates were then incubated at 56 °C for 5 min, followed by 98 °C for 5 min. Samples were diluted 1:10 in water and qPCR was performed with 3 μL of each diluted cell lysate using the LightCycler 480 Probes Master mix (Roche), according to the manufacturer’s instructions. Probes and primers used are as described: For *Wolbachia* detection, *16S rRNA* F (5’-GAGTGAAGAAGG CCTTTGGG-3’), *16S rRNA* R (5’-CACGGAGTTAGCCAGGACTTC-3’), and *16S rRNA* Cy5 probe (5’-LC640CTGTGAGTACCGTCATTATCTTCCTCACT-IowaBlackRQ-3’) were used. To detect the housekeeping gene, *RPS17* F (5′-TCCGTGGTATCTCCATCAAGCT-3′), *RPS17* R (5′-CACTTCCGGCACGTAGTTGTC-3′), and *RPS17* FAM probe (5’FAM-CAGGAGGAGGAACGTGAGCGCAG-BHQ1-3’) were utilized. PCR cycling conditions were 95 °C for 5 minutes, 45 cycles of 95 °C for 10 seconds, 60 °C for 15 seconds, 72 °C for 1 second, followed by cooling at 40 °C for 10 seconds. *Wolbachia* densities were quantified using the delta CT method (2^CT(reference)^/ 2^CT^(target)) ^94^.

### DENV-2

Dengue virus serotype 2 (DENV-2) strain 92T (isolated from human serum collected from a patient from Townsville, Queensland/Australia, in 1992) ^13^ was prepared by inoculation of C6/36 cell lines with a multiplicity of infection (MOI) of 0.1 and collection of culture supernatant 13-14 days post-infection. Infectious titers were determined by TCID_50_.

### DENV-2 infection of cell lines

*Aag2* cell lines were seeded to reach 90% confluency in 24, 12 or 6 well-plates and incubated at 26°C for 48 hours prior to viral infection. Unless otherwise stated, cells were infected with DENV-2 at MOI 1 in maintenance medium without FBS for 2 hours at 26°C. Virus inoculum was then removed and the cells washed once with warm PBS. Maintenance media containing 2% FBS was added and cells were then incubated at 26°C for the indicated time.

### Quantification of DENV-2 RNA

To quantify DENV-2 genomic copies, total RNA from virus-infected cells was extracted using the RNeasy kit (Qiagen) or Isolate II RNA mini kit (Bioline). DENV-2 RNA was amplified by qRT-PCR (LightCycler Multiplex RNA Virus Master, Roche), using primers to the conserved 3’UTR: Forward 5’-AAGGACTAGAGGTTAGAGGAGACCC; Reverse 5’-CGTTCTGTGCC TGGAATGATG; Probe 5’-HEX-AACAGCATATTGACGCTGGGAGAGACCAGA-BHQ13’ ^23^; DENV-2 RNA copies were quantified relative to *Ae. aegypti* house-keeping gene *RPS17* (primers and probe sequences as above) using the delta CT method (2^CT(reference)^/ 2^CT(target)^) ^94^. Reactions were run on a LightCycler 480 (Roche), and data analysis was carried out with the LightCycler 480 software. qRT-PCR conditions: 50 °C for 10 min, 95 °C for 30 s, followed by 45 cycles of 95 °C for 10 s, 55 °C for 5 s, and 72 °C for 10 s.

### TCID_50_ ELISAs to determine viral titres

Infectious virus levels were determined by TCID_50_ as previously described ^95^. Briefly, serial dilutions (10-fold) of virus were inoculated onto C6/36 cell lines in flat-bottom 96 well plates containing maintenance medium (RPMI-1640, 2% FBS), and incubated for 13-14 days at 28 °C. After the incubation period, media was aspirated, and cells were fixed with acetone fixative buffer (20% acetone/0.02% BSA in PBS) at 4 °C for 24 hours. The fixative was removed and the plate was left to completely air-dry. Once dried, virus concentrations were immediately quantified by ELISA or plates were kept at –20°C until required.

ELISAs were performed using monoclonal antibody 4G2 to detect DENV E protein_96_. Briefly, plates were blocked with 2% casein in TNE buffer (10mM Tris; 0.2M NaCl; 1mM EDTA; 0.05% Tween20) for 1 hour at room temperature. 4G2 (produced as hybridoma supernatant) ^97^ was diluted 1:200 in blocking solution and incubated on fixed cells for 1 hour at 37°C. Plates were washed four times with PBST (0.05% Tween20). Anti-mouse secondary antibody was diluted 1:2000 in blocking solution and incubated for 1 hour at 37°C. Plates were washed six times with PBST, then incubated with 3,3’5,5’-Tetramethylbenzidine (TMB) for up to 5 minutes. The reaction was stopped by addition of 0.1M HCl. The absorbance of each well was measured at 450nm using a BioTek Gen5 microplate reader (Agilent, Santa Clara, CA, USA). The TCID_50_/ml was determined using the Reed-Muench calculation ^98^.

### Cell fixation and processing for transmission electron microscopy

*Aag2* cell lines were seeded in a 6-well cell culture plate in triplicate at a concentration of 3 x 10^6^ cells/well for 48 hours at 26 °C. Unless otherwise stated, cells were mock-infected or infected for 2 hours at 26°C with DENV-2 at MOI 2 in maintenance media without FBS. Virus inoculum was then removed and cells were washed once with warm PBS. Maintenance media containing 2% FBS was added and cells were incubated at 26 °C for 2 days. For C75 treatment, cells were treated as described below and infected with DENV-2 at MOI 1. At 24 hours post-infection, cells were washed once with warm PBS and fixed for 2 hours at room temperature with 2% glutaraldehyde in 0.1 M sodium cacodylate buffer. Cells were gently washed three times for ten minutes each with 0.1 M sodium cacodylate buffer.

Cells were post fixed in 1% osmium tetroxide in 0.1M sodium cacodylate buffer for 30 minutes at room temperature which was then reduced with 1.5% potassium ferrocyanide in 0.1M sodium cacodylate buffer for 30 minutes. Cells were washed three times for 10 minutes with ultrapure water. Staining with 2.5% uranyl acetate (aqueous) followed overnight at 4°C. Cells were once again washed three times for 10 minutes with ultrapure water. Following washing, cells were scraped and pelleted into 4% agarose in water. Pellets were dehydrated with increasing concentrations of ethanol (25%, 50%, 75%, 90%, 100%, 100%) followed by acetone (100%, 100%). Each dehydration step was aided by a microwave regime of 40 seconds at 150 watts (Pelco Biowave). Pellets were infiltrated with epon resin using a microwave regime (25%, 50%, 75%, 100%, 100%; each step 3 minutes at 250 watts under vacuum). Pellets were transfer to fresh 100% epon and left at room temperature overnight and finally embedded in a silicone mold for polymersation for 48 hours at 60°C.

### Transmission electron microscopy

70nm sections were cut on a ultramicrotome (Leica UC7) and collected on copper mesh grids. Transmission electron microscopy (TEM) imaging was conducted on a Jeol JEM1400-Plus at 80kV. A combination of single snapshots and image montages were acquired, montages were automatically stitched by Jeol acquisition software (TEM Centre).

### Electron tomography

200nm sections were collected on copper mesh grids for electron tomography. 10nm gold fiducials were added to both sides of the section. Single axis tomography was performed on a Jeol JEM1400-Plus at 120kV using Jeol Recorder software. Tilt series were recorded with tilt angles from + 65° to − 65° with varying tilt increments based on a Saxton scheme. IMOD software ^99^ was used to reconstruct tilt series (etomo package), manual segmentation and visualisation.

### Immunofluorescence analyses

#### Live experiments

Cells were plated on 18 mm x 18 mm coverslips in a 6-well cell culture plate 48 hours previously coated with gelatin (0.2% [v/v]) ^63,84^. To stain the ER, the cell culture medium was aspirated, cells washed with PBS once and incubated for 30 minutes at 26°C with 1 μM live ER-tracker red dye (Molecular Probes) diluted in the appropriate *Aag2* cell culture medium. The ER-tracker solution was replaced by a 1/20 000 solution of SYTO-11 (Molecular Probes) DNA dye for 10 minutes at 26°C diluted in the appropriate *Aag2* cell culture medium, washed twice with PBS and cells were maintained in *Aag2* culture medium during the confocal microscopy observations.

#### LDs staining and quantification

Cells were plated on 12 mm x 12 mm coverslips in a 12-well cell culture plate for 48 hours previously coated with gelatin (0.2% [v/v]) ^63,84^. All cells were maintained in *Aag2* cell culture medium with 10% FBS. 24 hours post-DENV-2-infection, mock-infected and infected cells were fixed with 4% paraformaldehyde in PBS for 15 minutes, washed twice with PBS and permeabilized with 0.5% Saponin in PBS for 10 minutes and washed three times with PBS. Cells were blocked with 5% BSA for 1 hour, before antibody staining with polyclonal anti-WSP ^96^ and 4G2 hybridoma fluid against flavivirus group E antigen for DENV-2 staining. Cells were then incubated with Alexa Fluor 555 or 647 secondary antibodies (1:2000) for 1 hour. LDs were stained by incubating cells with BODIPY (493/503 4,4-difluoro 1,3,5,7,8 pentamethyl 4-bora3a,4a-diaza-s-indacene – Molecular Probes) at 1 ng/mL for 1 hour and nuclei were stained with DAPI (Sigma-Aldrich, 1 µg/ml) for 5 min. All incubations were at room temperature. Samples were then washed with PBS and mounted with Vectashield Antifade Mounting Medium (Vector Laboratories). The slides were imaged as 3-dimensional z-stacks and 2D images generated by Maximum Intensity Projection (MIP). For each condition, at least 6 fields of view were imaged at 63X magnification from different locations across each coverslip. LDs from at least 80 cells per biological replicate with a minimum of 2 biological replicates per experiment being analyzed for both LD number and average LD size. LD numbers and diameters were analyzed using quantitative data from the single raw CZI images (from Zen Blue) in ImageJ using the particle analysis tool.

For imaging of mosquito tissues, ovaries were dissected from non-blood-fed female mosquitoes 5-7 days post-emergence. Mosquito lines Rockefeller, Rockefeller-*w*Mel, Rockefeller-*w*AlbB and Rockefeller-*w*Pip have been described previously ^23^. Ovaries were dissected from 4-5 mosquitoes per line, in PBS, then fixed on poly-lysine coated slides in cold 4% paraformaldehyde/PBS for 15 minutes. Slides were rinsed three times in PBS, then permeabilized in 0.1% Triton X-100 in PBS for 10 minutes. Samples were blocked with 1% BSA for 30 minutes then stained with BODIPY (493/503) at 1 ng/ml for 1hour. Nuclei were stained with DAPI (Sigma-Aldrich, 1 μg/ml) for 5 minutes at room temperature. Samples were washed with PBS and mounted with Vectashield Antifade Mounting Medium (Vector Laboratories). All images were acquired using a Zeiss LSM 800 confocal microscope and processed using Zeiss analysis software (Zen Blue Edition version 10.1.19043, Jena, Germany) and FIJI analysis software. The slides were imaged as 3-dimensional z-stacks and 2D images generated by MIP.

### Inhibition of FAS by C75 treatment

Unless otherwise stated, *Wolbachia*-free and *w*Mel-*Aag2* lines were seeded and infected as described above. Prior to DENV-2 infection cells were pre-treated with 1 µM fatty acid synthase (FAS) inhibitor C75 (Abcam) or DMSO diluent only for 24 hours. Cells were mock-infected or DENV-2-infected at MOI 1 for 24 hours and C75 was maintained during the time of infection in *Aag2* cell culture medium with 10% FBS. LD quantification, viral replication and TEM imaging were performed as described above.

### Statistical analyses

Statistical analyses of parametric data were performed using One-way ANOVA or Two-way ANOVA tests with a Tukey’s multiple comparison correction, or unpaired two-tailed Student’s *t* test, and data were expressed as mean ± Standard Error of the Mean (±SEM). Statistical analyses of non-parametric data were performed using a Kruskal-Wallis test with Dunn’s correction for multiple comparisons, or a two-tailed Mann-Whitney test, and data were expressed as median ± Interquartile Range (± IQR). All statistical analyses were performed using Prism 9.3.1 (GraphPad Software), with p < 0.05 considered to be significant.

## Acknowledgements

This research was funded by the Australian Research Council (DP220102997) and the National Health and Medical Research Council (APP1182432). The funders had no role in study design, data collection and interpretation, or the decision to submit the work for publication.

We thank Professor Roy Hall and Jody Hobson-Peters (University of Queensland) for provision of the 4G2 antibody. The authors would like to acknowledge the Ramaciotti Centre for Cryo Electron Microscopy, a node of Microscopy Australia. The authors acknowledge Monash Micro Imaging, Monash University, for the provision of instrumentation, training, and technical support. We would like to thank the La Trobe University Bioimaging Facility for their assistance with image acquisition, data collection, and image analysis essential to this work. Finally, we thank Katrina Ibay for technical support.

## Author contributions

R.K.L performed the majority of the experiments; E.A.M assisted in the design of LD experiments, LD imaging and image analyses. R.T. and G.R. assisted with the TEM acquisition. E.A.M and R.T. equally collaborated to this work. J.T.B. originally established all mosquito cell lines and H.F. kindly provided the mosquito lines used in this study. J.M., K.J.H. and C.P.S. assisted in experimental direction. J.E.F. was responsible for the overall study design, alongside R.K.L. J.E.F. and C.P.S. acquired the funding for the project. J.E.F. and R.K.L. wrote the manuscript; all authors contributed to editing of the manuscript.

## Supplementary information for

## Supplementary text information

### Extended Material and Methods Mosquito rearing

All *Ae*. *aegypti* mosquitoes were reared and maintained as described previously ^1^. Briefly, adult mosquitoes were maintained at 26°C, 65% relative humidity, and a 12 h light:dark cycle in a climate-controlled room, and were allowed access to 10% sucrose *ad libitum.* Mosquitoes were blood fed on the arms of human volunteers for reproduction. To generate a panel of genetically comparable *Wolbachia*-carrying *Ae*. *aegypti* lines, we backcrossed females from the *w*Mel, *w*AlbB, and *w*Pip to males of the inbred laboratory *Ae*. *aegypti* line, Rockefeller ^2^ (BEI resources), for six generations. To exclude any influence of mosquito age on our experiments, age-controlled adults emerging within a 48 hours window were used.

### Live Lipid Droplet Staining

For live staining of LDs, *Wolbachia*-free and *w*Mel-*Aag2* cell lines were seeded in a 96-well black cell culture plate with 6 replicates per condition and at 1 x 10^5^ cells/well. Cells were incubated for 24 hours at 26 °C with *Aag2* cell culture medium with 10% FBS. After 24 hours, the culture medium was replaced with fresh medium containing DMSO only, 1 µM C75, 2.5 µM DGAT 1 (T863), 10 µM DGAT 2 (PF-06424439) or DGAT 1 and 2 combined, and incubated for another 24 hours. After incubation time, cell culture medium was replaced with 5 µM BODIPY 493/503 in cell culture medium for 1 hour at 26 °C. For each condition, extra wells were designated to not have cells or cells without BODIPY staining for appropriate background reduction. Cells were then gently washed 2 times with warm PBS and cells were maintained in PBS for fluorescence acquisition. Fluorescence was detected with a microplate reader (BioTek Gen5 - Agilent, Santa Clara, CA, USA) equipped for excitation in the 485/20 nm range and emission detection at 535/20 nm (FITC wavelength).

**Supplementary Fig. 1:**
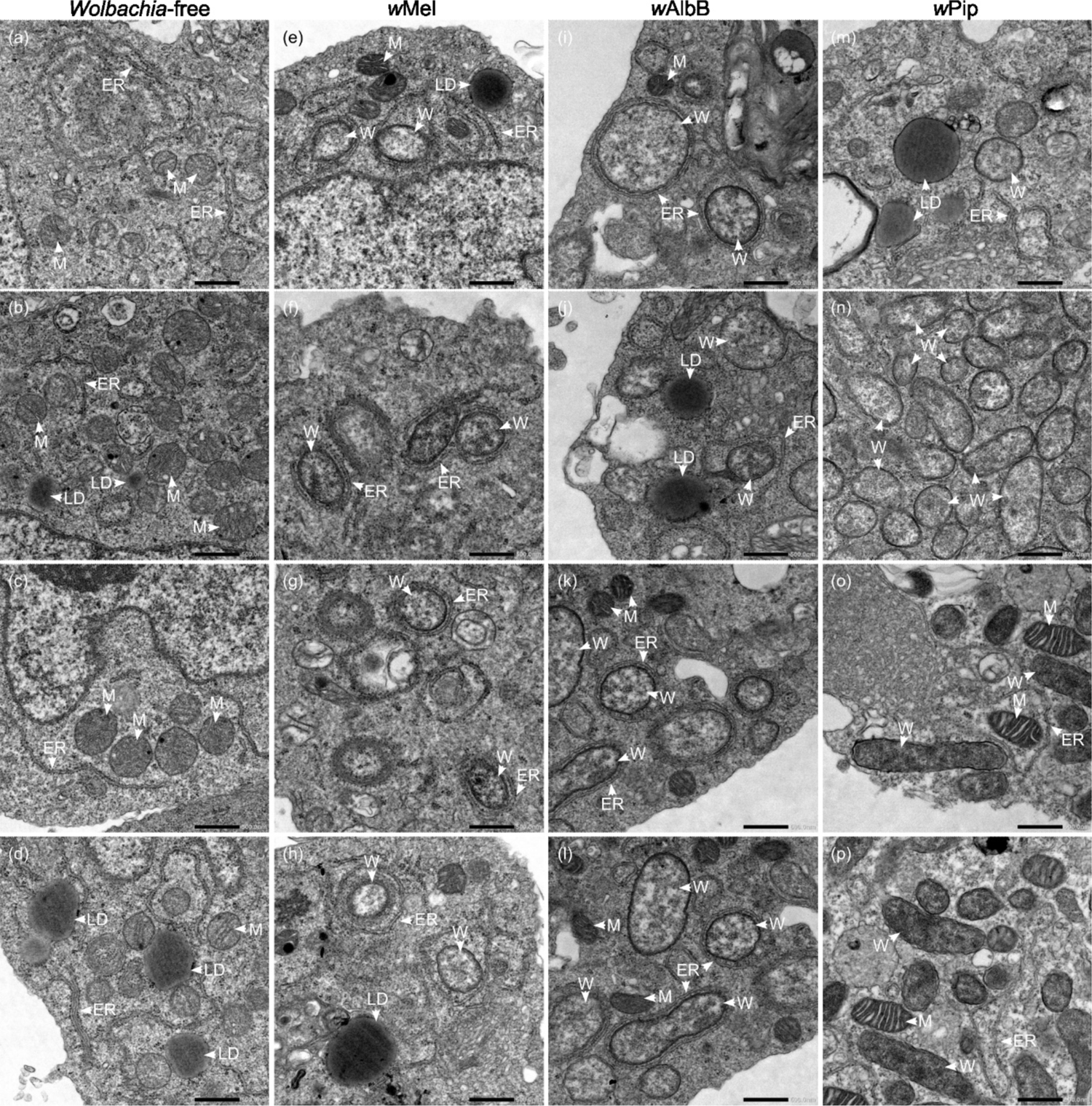
Antiviral *Wolbachia* strains are frequently wrapped by the host endoplasmic reticulum membranes. TEM micrographs of *Wolbachia*-free-*Aag2* cell line (a-d) or stably infected with *w*Mel (e-h), *w*AlbB (i-l), and *w*Pip (m-p) show their intracellular distribution and association with the ER membranes. Scale bar = 500nm. ER-Endoplasmic reticulum. W-*Wolbachia*. M-Mitochondria. LD-Lipid Droplet.

**Supplementary Fig. 2:**
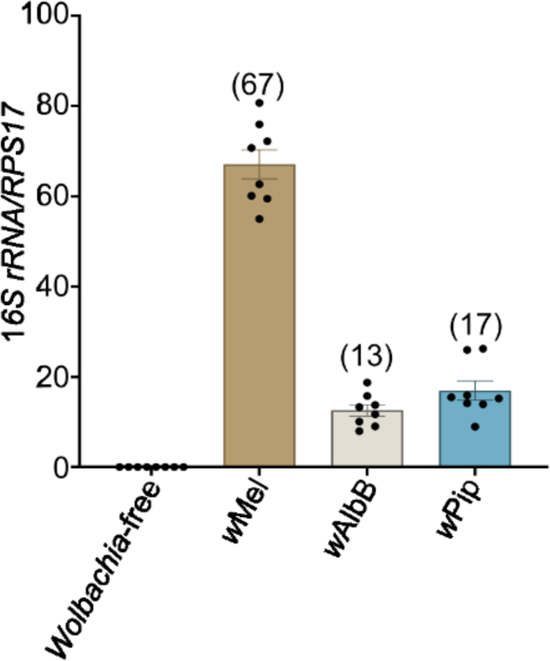
*Wolbachia* density in *Ae. aegypti* ovaries. Ovaries of at least 8 non-blood-fed female mosquitoes of Rockefeller (*Wolbachia*-free), Rockefeller-*w*Mel, Rockefeller-*w*AlbB, and Rockefeller-*w*Pip mosquito lines were dissected. *Wolbachia* per cell (in parentheses) was determined by qPCR (*Wolbachia 16S rRNA/RPS17*). Data are mean with ± Standard Error of the Mean (±SEM).

**Supplementary Fig. 3:**
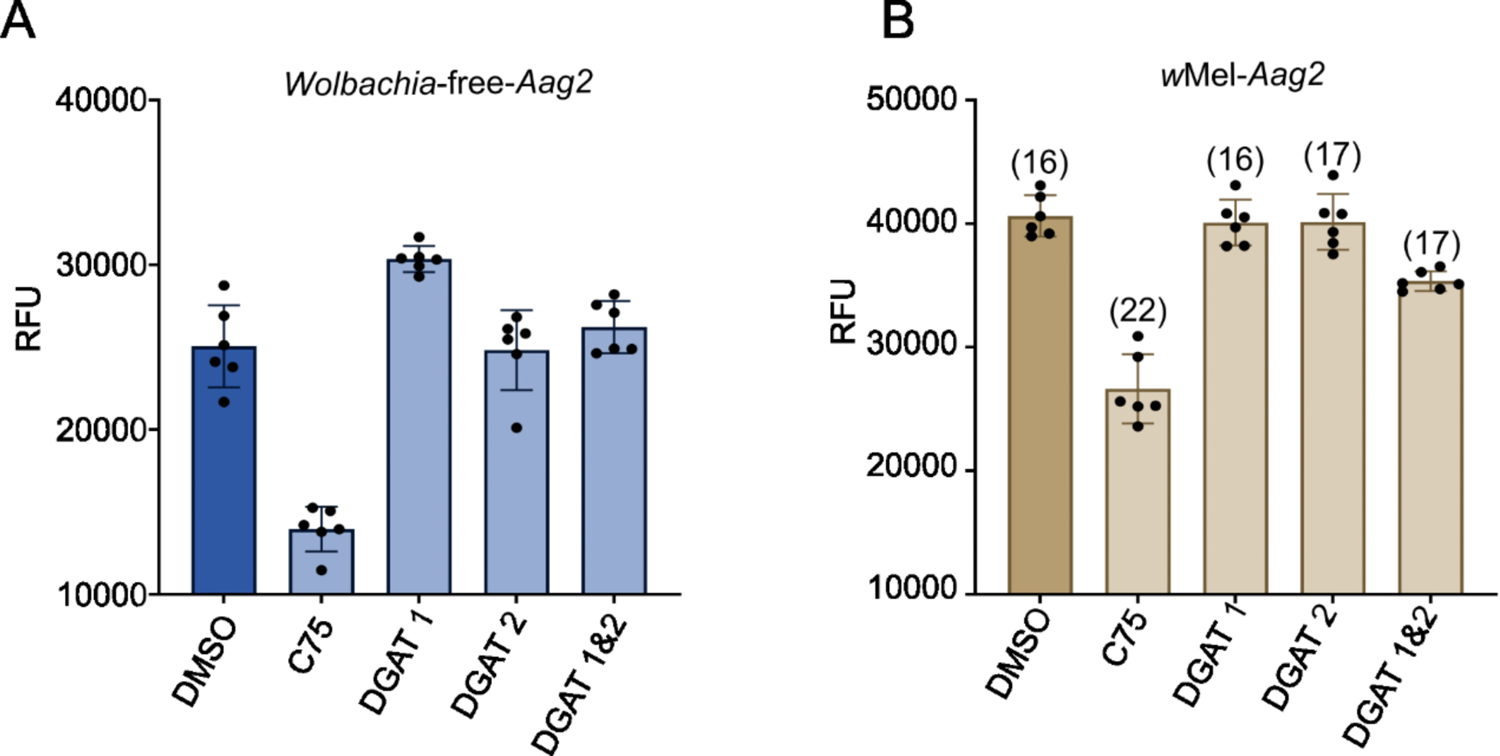
The Fatty Acid Synthase (FAS) inhibitor, C75, efficiently reduces LD accumulation in *Aag2* cell lines. (A) *Wolbachia*-free and (B) *w*Mel-*Aag2* cell lines were treated with C75, DGAT 1, DGAT 2, and DGAT 1&2 combined for 24 hours prior to LD staining. For live staining of LDs, cells were incubated with BODIPY 493/503 for 1 hour. The numbers in parentheses represent the average number of *Wolbachia* per cell (*Wolbachia 16S rRNA*/*RPS17*) after each treatment. Data are mean with ± Standard Deviation (SD) and represent 1 independent experiment with six biological replicates each. Fluorescence was detected with a microplate reader at 485/535 nm (excitation/emission) range. RFU – Relative fluorescence units.

