## Supplementary information for "Antiviral *Wolbachia* strains associate with *Aedes aegypti* endoplasmic reticulum membranes and disturb host cell lipid distribution to restrict dengue virus replication"

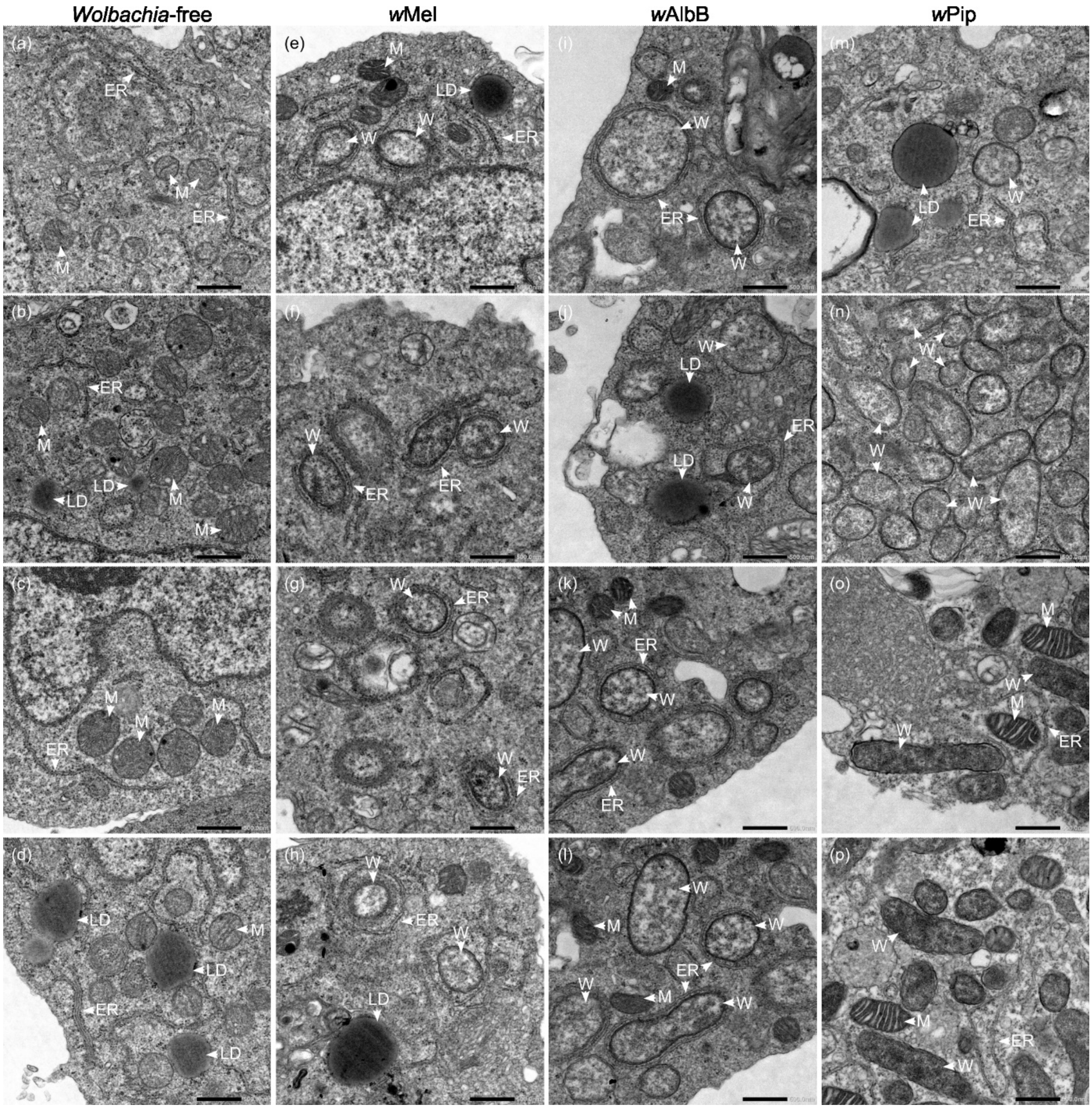


**Supplementary Fig. 1: Antiviral *Wolbachia* strains are frequently wrapped by the host endoplasmic reticulum membranes.** TEM micrographs of *Wolbachia*-free-*Aag2* cell line (a-d) or stably infected with *w*Mel (e-h), *w*AlbB (i-l), and *w*Pip (m-p) show their intracellular distribution and association with the ER membranes. Scale bar = 500nm. ER- Endoplasmic reticulum. W- *Wolbachia*. M- Mitochondria. LD- Lipid Droplet.

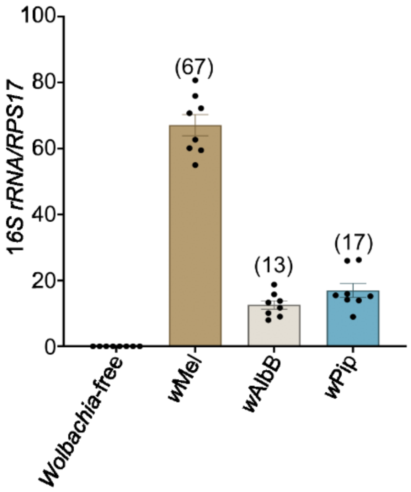


**Supplementary Fig. 2: *Wolbachia* density in *Ae. aegypti* ovaries.** Ovaries of at least 8 non-blood-fed female mosquitoes of Rockefeller (*Wolbachia*-free), Rockefeller-*w*Mel, Rockefeller-*w*AlbB, and Rockefeller-*w*Pip mosquito lines were dissected. *Wolbachia* per cell (in parentheses) was determined by qPCR (*Wolbachia 16S rRNA/RPS17*). Data are mean with ± Standard Error of the Mean (±SEM).

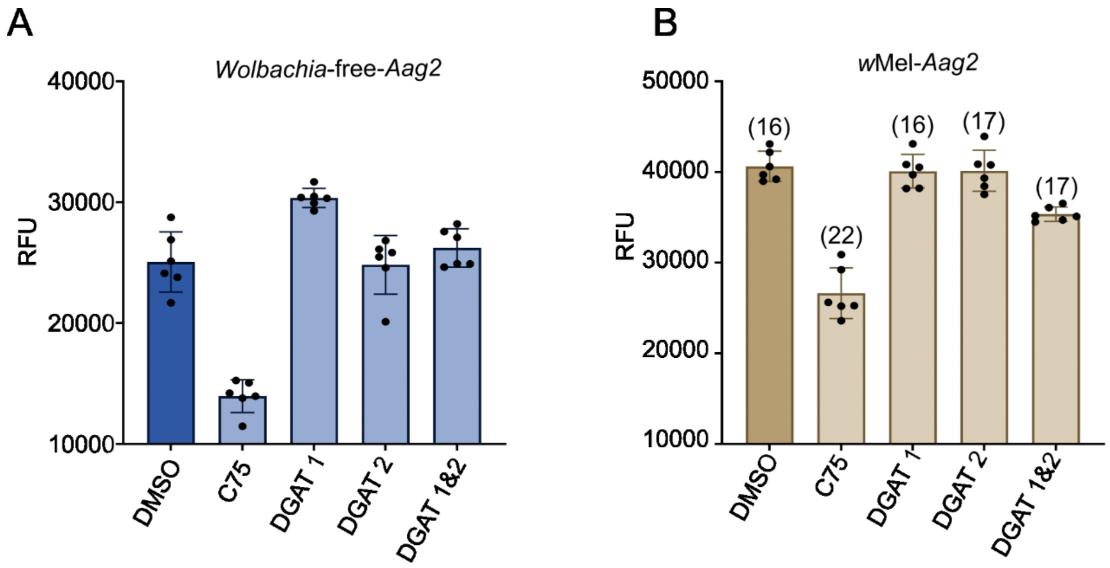


**Supplementary Fig. 3: The Fatty Acid Synthase (FAS) inhibitor, C75, efficiently reduces LD accumulation in *Aag2* cell lines.** (A) *Wolbachia*-free and (B) *w*Mel-*Aag2* cell lines were treated with C75, DGAT 1, DGAT 2, and DGAT 1&2 combined for 24 hours prior to LD staining. For live staining of LDs, cells were incubated with BODIPY 493/503 for 1 hour. The numbers in parentheses represent the average number of *Wolbachia* per cell (*Wolbachia 16S rRNA*/*RPS17*) after each treatment. Data are mean with ± Standard Deviation (SD) and represent 1 independent experiment with six biological replicates each. Fluorescence was detected with a microplate reader at 485/535 nm (excitation/emission) range. RFU – Relative fluorescence units.
